# Identification of pituitary thyrotrope signature genes and regulatory elements

**DOI:** 10.1101/2020.08.05.238253

**Authors:** Alexandre Z. Daly, Lindsey A. Dudley, Michael T. Peel, Stephen A. Liebhaber, Stephen C. J. Parker, Sally A. Camper

## Abstract

Pituitary thyrotropes are specialized cells that produce thyroid stimulating hormone (TSH), a critical factor for growth and maintenance of metabolism. The transcription factors POU1F1 and GATA2 have been implicated in thyrotrope fate and transcriptional regulation of the beta subunit of TSH, *Tshb*, but no transcriptomic or epigenomic analyses of these cells has been undertaken. The goal of this work was to discover key transcriptional regulatory elements that drive thyrotrope fate. We identified the transcription factors and epigenomic changes in chromatin that are associated with differentiation of POU1F1-expressing progenitors into thyrotropes, a process modeled by two cell lines: one that represents an early, undifferentiated *Pou1f1* lineage progenitor (GHF-T1) and one that is a committed thyrotrope that produces TSH (TαT1). We generated and compared RNA-seq, ATAC-seq, histone modification (including H3K27Ac, H3K4Me1, and H3K27Me3), and transcription factor (POU1F1) binding in these two cell lines to identify regulatory elements and candidate transcriptional regulators. We identified POU1F1 binding sites that were unique to each cell line. POU1F1 binding sites are commonly associated with bZIP transcription factor consensus binding sites in GHF-T1 cells and Helix-Turn-Helix (HTH) or basic Helix-Loop-Helix (bHLH) factors in TαT1 cells, suggesting that these classes of transcription factors may recruit or cooperate with POU1F1 binding to unique sites. We validated enhancer function of novel elements we mapped near *Cga, Pitx1, Gata2,* and *Tshb* by transfection in TαT1 cells. Finally, we confirmed that an enhancer element near *Tshb* can drive expression in thyrotropes of transgenic mice, and we demonstrate that GATA2 enhances *Tshb* expression through this element. These results extend the ENCODE multi-omic profiling approach to an organ that is critical for growth and metabolism, which should be valuable for understanding pituitary development and disease pathogenesis.

## Introduction

Recent genome-wide association studies (GWAS) have begun to identify loci that are associated with sporadic pituitary adenomas and variation in normal height, but the genes associated with many of these loci are unknown (Jarvis et al. 2012; Turchin et al. 2012; Ye et al. 2015). Nearly 90% of GWAS hits are in noncoding regions, making it difficult to transition from genetic mapping to biological mechanism (Edwards et al. 2013). Recent studies that identify enhancer regions by undertaking large scale functional genomic annotation of non-coding elements like Encyclopedia of DNA Elements (ENCODE) have begun to yield a better understanding of some complex diseases. Dense molecular profiling maps of the transcriptome and epigenome have been generated for more than 250 cell lines and 150 tissues, but pituitary cell lines or tissues were not included. This represents a major limitation, as the cell types that comprise the pituitary gland secrete hormones responsible for growth (growth hormone secreted by somatotropes), reproduction (gonadotropins secreted by gonadotropes), adrenal gland function and the stress response (ACTH secreted by corticotropes), lactation (prolactin secreted by lactotropes), and thyroid gland function (thyroid-stimulating hormone secreted by thyrotropes). Epigenomic and gene expression data are emerging for somatotropes, gonadotropes and corticotropes, but there is very little available data on thyrotropes (Qiao et al. 2016; Mayran et al. 2018; Peel et al. 2018; Mayran et al. 2019).

Thyrotropes represent ~5% of cells in the pituitary gland, and their function is to express and secrete Thyroid Stimulating Hormone (TSH or thyrotropin), which regulates thyroid gland development and thyroid hormone production. These hormones are essential for normal growth and metabolism. Up to 12% of the US population suffers from abnormal levels of thyrotropin (Canaris et al. 2000). The incidence of secondary hypothyroidism is estimated to be 1:20,000 to 1:80,000 individuals (Persani et al. 2019). Research into the regulation of thyrotrope differentiation and function is relevant to this public health problem.

A cascade of transcription factors is responsible for the differentiation of the major pituitary hormone-producing cell types, and three transcription factors associated with thyrotrope development and function are POU1F1, GATA2, and ISL1.The pituitary transcription factor POU1F1 is essential for the differentiation of growth hormone, prolactin and TSH-producing cells (Li et al. 1990). It binds to the promoters of *Gh, Prl,* and *Tshb* to activate gene expression (Bodner and Karin 1987; Ingraham et al. 1988; Gordon et al. 1997; Wood et al. 1999; Hashimoto et al. 2000). Defects in the POU1F1 gene cause severe growth insufficiency and hypothyroidism in humans and mice (Li et al. 1990; Fang et al. 2016). POU1F1 and GATA2 act synergistically to activate *Tshb* expression through promoter-proximal elements (Gordon et al. 1997; Dasen et al. 1999). Defects in GATA2 and ISL1 reduce thyrotrope differentiation in mice, but they do not appear to ablate it (Charles et al. 2006; Castinetti et al. 2015; Brinkmeier et al. 2020). Despite the important role of *Pou1f1* in thyrotrope development and function, little is known about the gene regulatory network of POU1F1 in progenitors or thyrotropes.

Due to the scarcity of the thyrotrope cell-type, classical genomic techniques are challenging to apply. Hormone-producing cell lines have been invaluable for understanding changes in chromatin and gene expression that occur during development (Budry et al. 2012; Mayran et al. 2018). To discover thyrotrope-specific regulatory elements and potential drivers of differentiation, we generated and compared the transcriptome (RNA-seq), open chromatin (ATAC-seq), histone modification (CUT&RUN for H3K27Ac, H3K4Me1, and H3K27Me3), and transcription factor binding (CUT&RUN for POU1F1) in two mouse cell lines, a POU1F1-expressing pituitary precursor cell line that does not express any hormones, GHF-T1, to a thyrotrope-like cell line, TαT1 (Lew et al. 1993; Alarid et al. 1996). TαT1 cells behave much like endogenous thyrotropes in that they respond to TRH and retinoids, and secrete TSH in response to diurnal cues (Janssen et al. 2011; Nakajima et al. 2012; Aninye et al. 2014). Finally, we evaluated putative enhancer elements for function using transfection assays in TαT1 cells and genetically engineered mice. Together, these studies extend ENCODE-like multi-omic analyses to generate reference maps of gene regulation for cell types critical for growth and metabolism.

## Results

### Comparison of transcriptomes

To identify candidate factors that drive the differentiation of thyrotropes, we performed RNA-sequencing on the GHF-T1 and TαT1 cell lines. There were many differences in their transcriptomes, consistent with their distinctive morphology, growth rate, and hormone secretion properties (**Fig. 1 A**). Eighty-two percent of genes were differentially expressed (FDR < 0.01). *Pou1f1* expression levels were nearly twice as high in TαT1 cells (160 FPKM) relative to GHF-T1 cells (85 FPKM). Other SV40 immortalized pituitary cell lines vary ten-fold in *Pou1f1* expression levels, but there was no correlation with differentiation state (Sizova et al. 2010). As expected, *Cga* and *Tshb* were expressed in TαT1 (3557 and 11 FPKM, respectively) but negligibly expressed in GHF-T1 cells (1.4 and 0 FPKM, respectively). The GHF-T1 cells had elevated expression of the transcription factors *Gli3*, *Pax3*, and *Foxg1* (**Table 1**). TαT1 cells had elevated expression of *Gata2* and *Isl1,* as expected, and *Lhx3*, *Rxrg*, *Neurod4,* and *Insm1* were also significantly up regulated.

**Figure 1:**
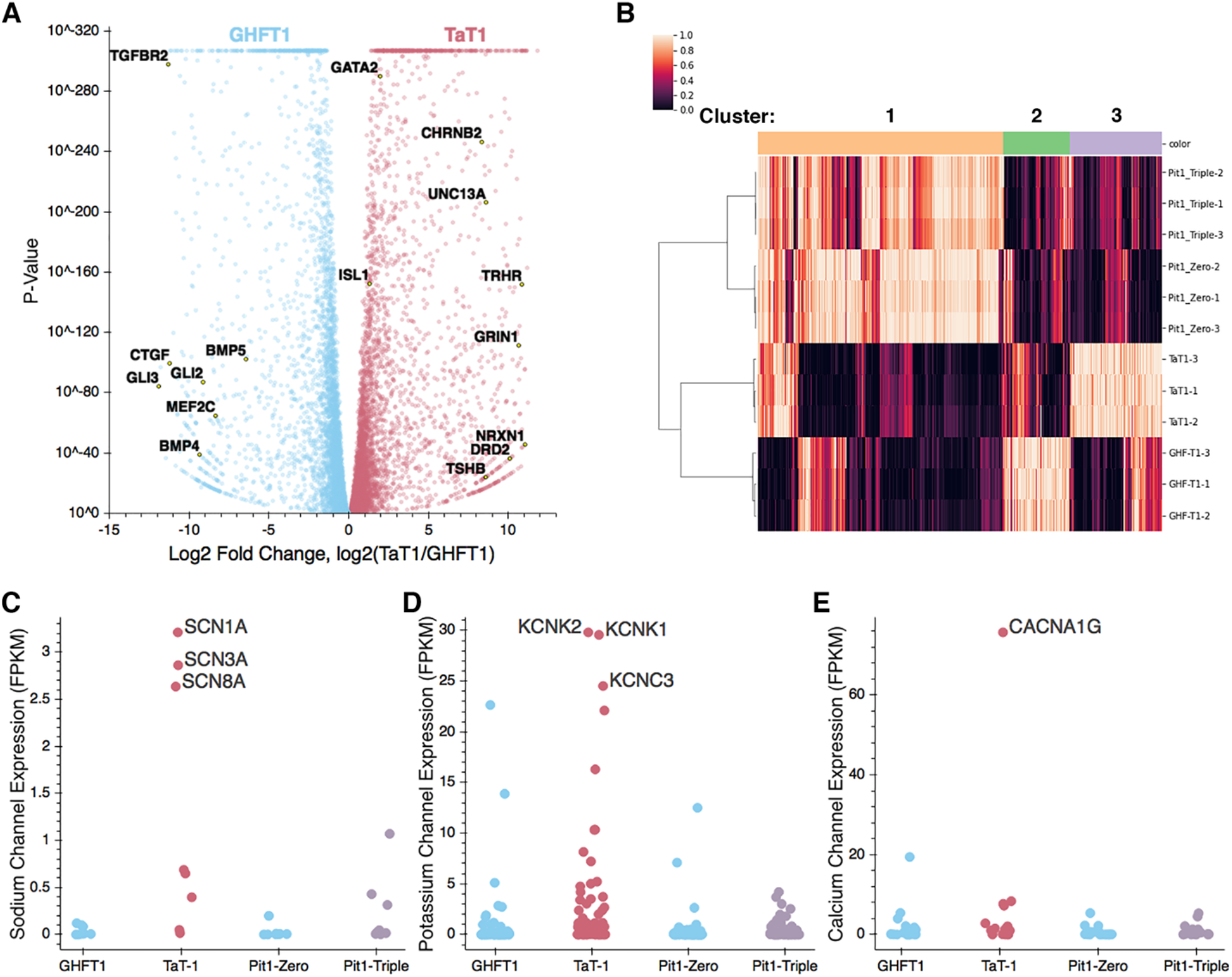
(**A**) Volcano plot of differential gene expression for GHF-T1 compared to TαT1 cells. Genes upregulated in GHF-T1 cells are colored blue, and those upregulated in TαT1 cells are colored red. Labeled genes represent key factors and genes associated with GO terms in **Table 2**. (**B**) Heatmap showing similarly and differentially expressed genes across GHF-T1, TαT1, Pit1-Zero and Pit1-Triple cells. Genes associated with each cluster are presented in **Supplemental Table 1**. (**C**) FPKM values of neuronal sodium channel genes. (**D**) FPKM values of potassium channel genes. (**E**) FPKM values of calcium channel genes.

**Table 1.** Differentially expressed transcription factors (FDR <5×10^−14^)

| Table 1. Differentially expressed transcription factors (FDR <5x10<sup>-14</sup>) |  |  |  |  |  |
| --- | --- | --- | --- | --- | --- |
| GHFT1 cells |  |  | TαT1 cells |  |  |
| Gene | Log2 fold change | rank | Gene | Log2 fold change | rank |
| *Gli3* | 11.90 | 1 | *Lhx3* | 11.04 | 8 |
| *Pax3* | 10.95 | 12 | *Rxrg* | 10.74 | 25 |
| *Zfhx4* | 10.74 | 21 | *Insm1* | 10.67 | 30 |
| *Foxg1* | 10.59 | 26 | *Neurod4* | 10.56 | 37 |
| *Zfp57* | 10.51 | 29 | *Sox3* | 10.23 | 55 |
| *Hoxc13* | 10.29 | 40 | *Fev* | 10.18 | 57 |
| *Kdm5D* | 10.19 | 48 | *Myt1l* | 9.93 | 72 |
| *Hmga2* | 10.15 | 53 | *Scrt2* | 9.90 | 77 |
| *Msx2* | 10.11 | 54 | *Zfp641* | 9.39 | 126 |
| *Runx1* | 10.07 | 55 | *Scrt1* | 9.15 | 145 |
| *Rhox10* | 9.94 | 60 | *Zic3* | 8.96 | 168 |
| *Mecom* | 9.84 | 68 | *En2* | 8.95 | 169 |
| *Tead2* | 9.65 | 80 | *Lin28b* | 8.82 | 184 |
| *Vax1* | 9.58 | 85 | *Pax5* | 8.61 | 205 |
| *Tcf7l1* | 9.57 | 87 | *Zfp709* | 8.52 | 218 |
| *Hoxa1* | 9.52 | 91 | *Prdm16* | 8.30 | 249 |
| *Tcf24* | 9.46 | 95 | *Pou2f2* | 8.15 | 268 |
| *Hoxc9* | 9.35 | 104 | *Zim1* | 7.92 | 301 |
| *Maf* | 9.32 | 106 | *Nhlh1* | 7.85 | 312 |
| *Pou3f3* | 9.16 | 112 | *Foxl2* | 7.82 | 317 |

To uncover pathways up- and down-regulated in these cell lines, we performed GO-term (gene ontology) enrichment analysis on the top 5% of the most differentially expressed genes (by log-2 fold-change) in both lines (Ashburner et al. 2000; The Gene Ontology 2019). The GO terms enriched in GHFT1 cells were broadly related to development and morphogenesis (**Table 2, Supplemental Fig. 1**). Genes contributing to these GO terms include genes from the GLI family that are targets of hedgehog signaling (*Gli2*, *Gli3*, *Glipr1*, *Glipr2*, *Glis2*, *Glis3*), BMPs (*Bmp1*, *Bmp4*, *Bmp5*, *Bmpr1a*, *Bmpr2*, *Bmper*, *Bmp2k*), and FGFs (*Fgf5*, *Fgf7*, *Fgf8*, *Fgf10*, and *Fgf21*). Increased expression of these factors in GHFT1 cells is consistent with the underlying the importance of FGF, BMP, and sonic hedgehog signaling in early pituitary development (Ericson et al. 1998; Treier et al. 2001; Carreno et al. 2017). In contrast, the up-regulated genes in TαT1 cells were enriched for GO terms related to nervous system development and synapse formation. Some genes enriched in TαT1 cells that contribute to these neuronal GO terms are neurexin, genes of the glutamate receptor family (*Grin1*, *Grina*, *Grin2d*, *Grin3a*), and the synaptic regulator *Unc13a*. KEGG-pathway enrichment analysis revealed an increase in neuroactive ligand-receptor interaction in TαT1 cells, consistent with the enrichment in GO terms found.

**Table 2.**
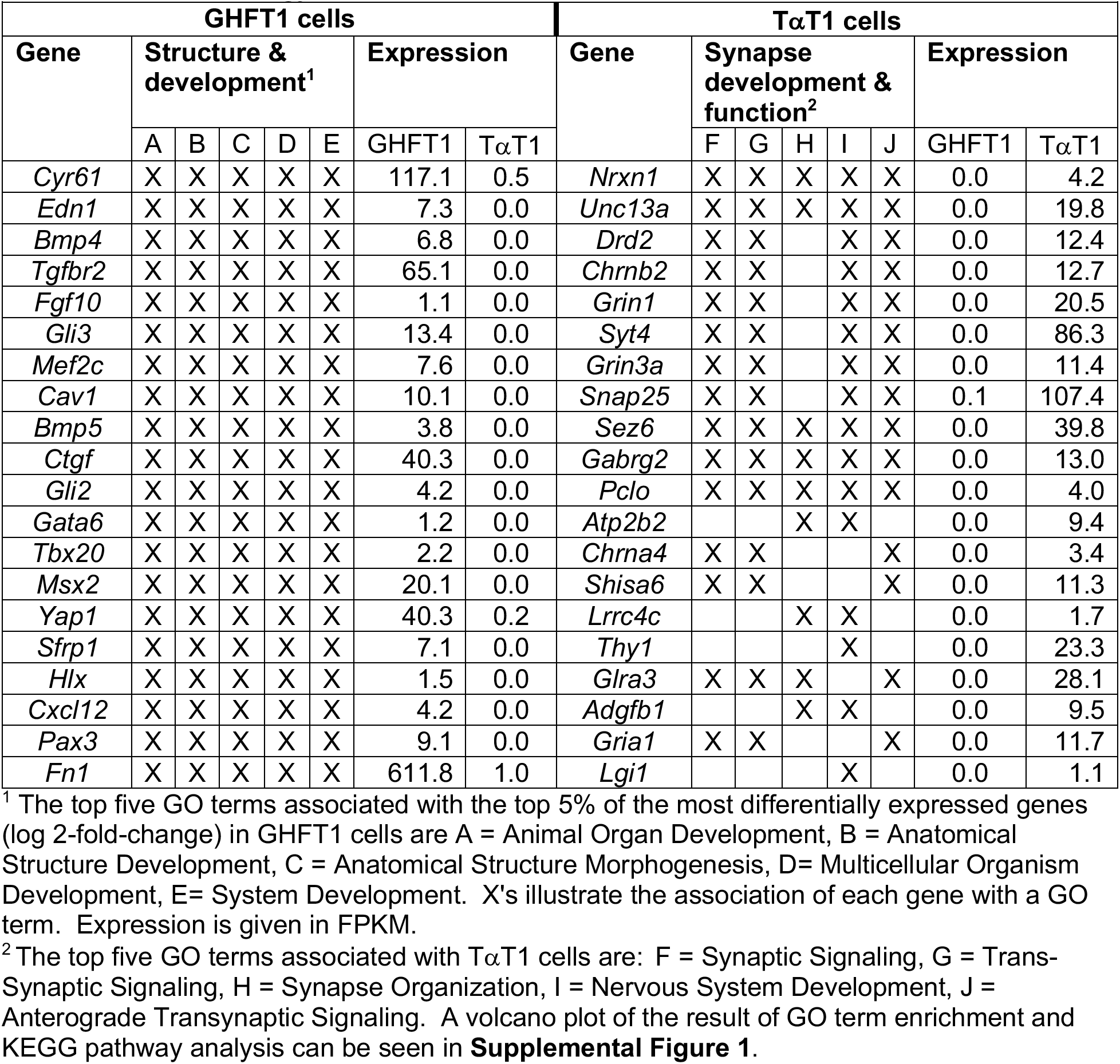
Gene ontology term enrichment.

The function of several members of the bHLH family of transcription factors, including *Ascl1, Neurod4,* and *Neurod1* has been investigated in pituitary development (Ando et al. 2018). Seventy-one of the ninety-three known bHLH factors are differentially expressed between GHF-T1 and TαT1 cells (FDR < 0.05); *Neurod4* and *Ascl1* are upregulated in TαT1 cells (**Supplemental Table 2**). *Ascl1* is essential for development of all hormone-producing cell types in fish pituitary, and in mice, *Ascl1* loss of function causes reduced production of *Pomc*, *Lhb*, and *Fshb* (Zhang et al. 2015; Ando et al. 2018). However, these reports conflict on whether thyrotropes are affected by *Ascl1* deficiency. We performed TSH immunostaining on pituitaries from *Ascl1*-null mice and did not detect a reduction in thyrotropes at e18.5 (**Supplemental Figure 2**), suggesting *Ascl1* is not required for thyrotrope cell specification. Repressive bHLH genes of the ID family had the highest expression in both of the cell lines, and the role of these genes has not been investigated.

We compared gene expression profiles that we obtained from GHF-T1 and TαT1 cells with those of other SV40-transformed pituitary cell lines, Pit1-zero and Pit1-triple cells (Sizova et al. 2010). Pit1-zero and Pit1-triple cells were transformed using the same *Pou1f1* regulatory elements as the GHF-T1 cell line. Pit1-zero cells express *Pou1f1*, but none of POU1F1’s downstream hormone genes, whereas Pit1-triple cells express *Pou1f1* and all three POU1F1-dependent hormones, GH, PRL, and TSH. In the TαT1 cell line, we found a statistically significant increase in the expression of sodium channel genes (p-value = 0.002) and potassium channel genes (p-value = 2.9e-05), but not in calcium channel genes as a group (p-value = 0.26). The most highly expressed sodium channel genes in TαT1 cells are *Scn1a*, *Scn8a*, and *Scn3a*. Three sodium channel genes (*Scn1a*, *Scn8a*, and *Scn9a*) are also expressed in the Pit1-Triple cell line, the only other hormone-expressing cell lineage we studied. The most highly expressed potassium channel genes in TαT1 cells are *Kcnc3*, *Kcnq2*, *Kcnk1*, and *Kcnk2*. G protein-gated ion channels are involved in regulated hormone secretion. This marked increase in ion channel genes in the TαT1 cells is consistent with their GO terms associated with synapses and neuron formation and function. The only calcium channel gene with differential expression was *Cacna1g,* which is highly expressed in TαT1 cells relative to the other three cell lines.

### Chromatin landscape around thyrotrope-signature genes

To assess genome-wide changes in the chromatin landscape associated with thyrotrope differentiation, we performed Cleavage Under Target and Release using Nuclease (CUT&RUN) for three major histone marks: H3K27Ac, H3K4Me1, and H3K27Me3 (Skene and Henikoff 2017). The presence of both H3K27Ac and H3K4Me1 mark active enhancers, while H3K27Me3 marks repressed regions (Boyer et al. 2006; Bracken et al. 2006; Lee et al. 2006; Creyghton et al. 2010). We also performed an Assay for Transposase-Accessible Chromatin with High-Throughput Sequencing (ATAC-seq), a method for profiling regions of accessible chromatin, which are often regulatory (Buenrostro et al. 2013). The results for three selected genes, *Isl1, Gli3,* and *Rxrg* and are shown in **Figure 2.** These data (called tracks) reveal that *Isl1* is expressed in both cell lines, and has extensive H3K27Ac, H3K4Me1, and ATAC-seq signal across the locus, revealing putative active enhancers and areas of open chromatin. For example, the stretch of H3K4Me1 and H3K27Ac signal covering the last intron and penultimate exon of *Isl1* could be an enhancer (**Fig. 2A**).

**Figure 2:**
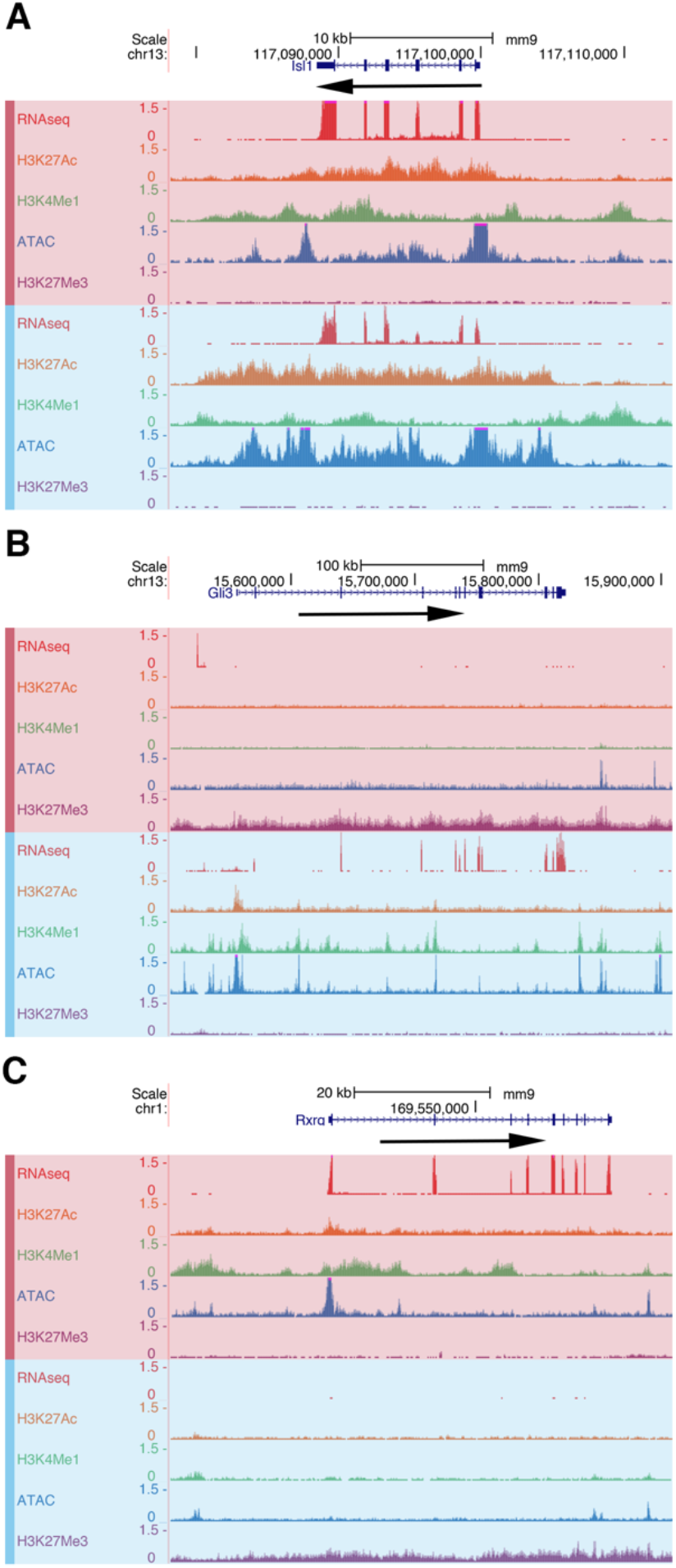
(**A**) *Isl1* encodes a key pituitary transcription factor that is expressed in TαT1 (red) and GHF-T1 (blue) cell lines. Data from RNA-Seq, CUT&RUN for active chromatin marks (H3K27Ac and H3K4Me1), open chromatin (ATAC-seq), and CUT&RUN for repressive chromatin marks (H3K27Me3) is visualized for each cell line in genome browser tracks. (**B**) *Gli3* is in active chromatin and expressed in GHF-T1 cells. *Gli3* is not expressed in TαT1 cells, and the chromatin is inaccessible with repressive marks. (**C**) *Rxrg* is expressed and in active chromatin in TαT1 cells, while not expressed and repressed in GHF-T1 cells.

We visualized the expression and chromatin architecture around genes that are differentially expressed in the precursor and differentiated cell lines. Here we show the tracks for *Gli3* and *Rxrg* which are the most and second-most differentially expressed transcription factors in the GHF-T1 and TαT1 cells, respectively. *Gli3* is strongly expressed in GHF-T1 cells, and has many H3K27Ac, H3K4Me1, and ATAC-seq peaks, revealing active enhancers in areas of open chromatin (**Fig. 2B**). By contrast, *Gli3* is not expressed in TαT1 cells. The chromatin surrounding *Gli3* in the TαT1 cells is devoid of H3K27Ac, H3K4Me1, and ATAC-seq peaks and is covered with H3K27Me3, a mark of active repression. This shows that *Gli3* is not expressed and is actively repressed in the TαT1 cell line. Conversely, *Rxrg*, a gene whose deletion in mice is associated with thyroid hormone resistance, is highly expressed in TαT1 cells but not in GHF-T1 cells (Brown et al. 2000). Consistent with this, in TαT1 cells the *Rxrg* locus is decorated with H3K27Ac, H3K4Me1, and ATAC-seq peaks, whereas the GHF-T1 line has no such peaks, and shows active repression of *Rxrg*, with a broad H3K27Me3 signal (**Fig. 2C**).

We used ChromHMM to annotate different chromatin states based on H3K27Ac, H3K4Me1, H3K27Me3, and ATAC-seq signal (Ernst and Kellis 2012). Iterating over increasing numbers of possible states, we found that 11 states best captured the chromatin architecture within these two cell lines (**Supplemental Fig. 6**). Of these states, two had both H3K4Me1 and H3K27Ac, indicating active enhancers. The difference between the two states was the presence or absence of an ATAC-seq signal, meaning one state represented open, active enhancers, while the other represented active enhancers in a more closed state.

### POU1F1 binding

We performed CUT&RUN for POU1F1 in both the GHF-T1 and TαT1 cell lines to identify similarities and differences in POU1F1 at these two stages of differentiation. *Pou1f1* has two enhancers, a proximal (5.6 kb), early-stage enhancer bound by PROP1, and a distal (10 kb), late-stage enhancer which POU1F1 binds to and drives its own expression in an auto-regulatory fashion after birth (Chen et al. 1990; DiMattia et al. 1997). In both cell lines CUT&RUN shows extensive POU1F1 binding across the *Pou1f1* promoter-proximal region and both the early and late enhancers (**Fig. 3A**). The *Twist1* promoter is an example of preferential POU1F1 binding in GHF-T1 cells relative to TαT1 cells (**Fig. 3B**). TWIST1 is a bHLH protein that plays an important role in head development and is mutated in patients with Saethre-Chotzen syndrome (Kress et al. 2006). *Twist1* is more highly expressed in GHF-T1 than TαT1 cells (having a log-2-fold change value of 5). Of note, POU1F1 binding was detected at the neurexin promoter in TαT1 cells, and neurexin expression increased from nearly zero in GHFT1 cells to 5 FPKM in TαT1 cells (**Fig. 3C**). Neurexin is critical for proper synapse formation.

**Figure 3:**
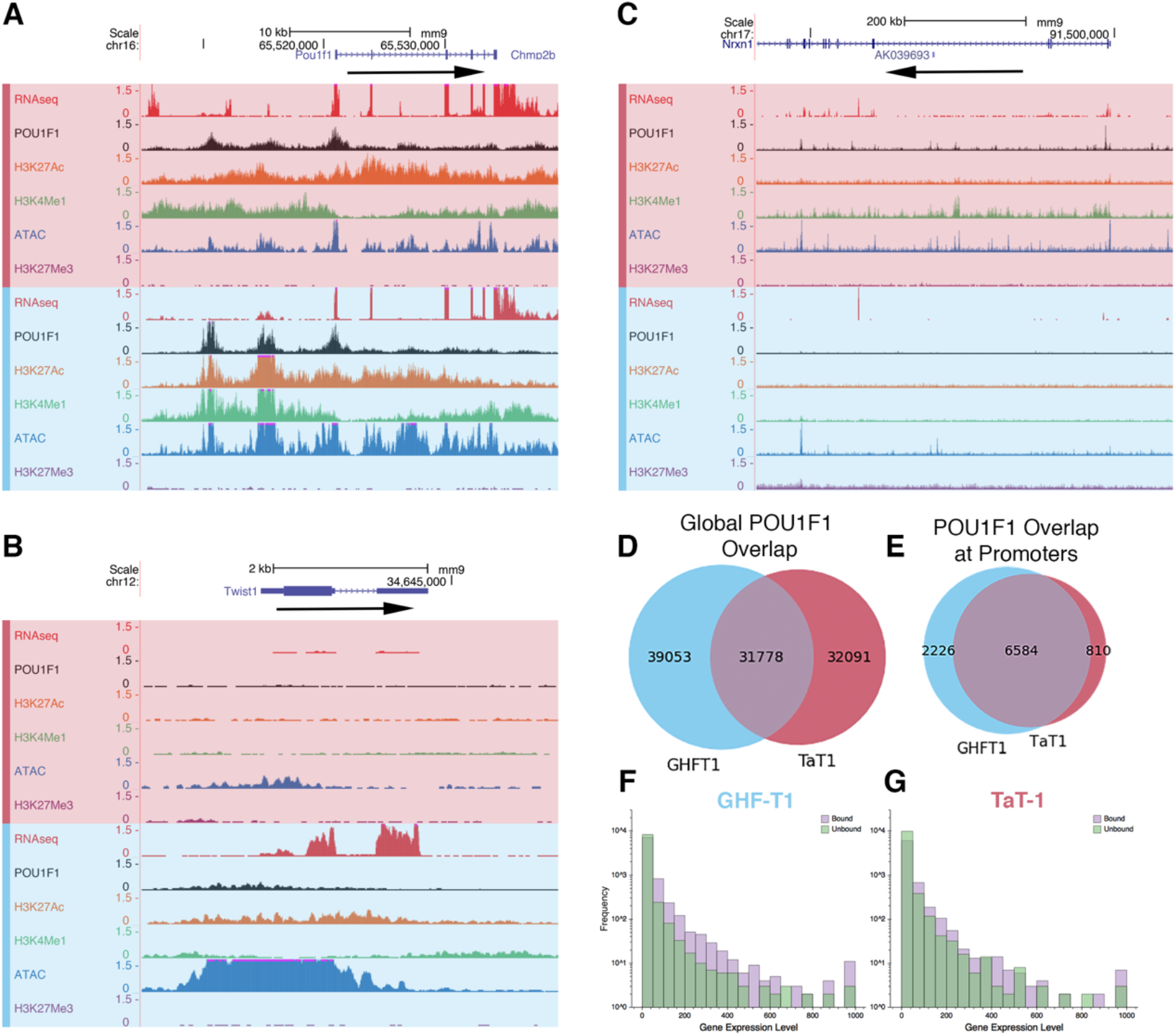
(**A**) *Pou1f1* is expressed in both cell lines, contains active chromatin marks, and exhibits similar POU1F1 binding to enhancer elements. Data from RNA-seq, CUT&RUN for POU1F1 and active chromatin marks (H3K27Ac, H3K4Me1), ATAC-seq, and CUT&RUN for repressive marks (H3K27Me3) is visualized in genome browser tracks for TαT1 cells (red) and GHF-T1 (blue). (**B**) *Twist1* is an example of a gene uniquely bound by POU1F1 and expressed in GHF-T1 cells. (**C**) *Nrxn1* is an example of a gene uniquely bound by POU1F1 and expressed in TαT1 cells. (**D**) A Venn diagram illustrating the number of shared and distinct POU1F1 binding sites throughout the genomes of GHF-T1 and TαT1 cells. (**E**) A Venn diagram showing the number of shared and distinct genes whose promoters are bound by POU1F1 in GHF-T1 and TαT1 cells. (**F**) A histogram showing expression of genes in GHF-T1 cells that have POU1F1 bound to their promoters in purple and that do not have POU1F1 bound to their promoters in green. (G) A histogram showing expression of genes in TαT1 cells that have POU1F1 bound to their promoters in purple and that do not have POU1F1 bound to their promoters in green.

Genome-wide analysis of POU1F1 binding at promoters revealed that only 15-16% of POU1F1 binding sites in GHF-T1 cells (10,980 out of 69,644) and TαT1 cells (9,360 out of 63,036) are within 1 kb of a transcription start site (TSS). While only one third of all POU1F1 binding sites are shared between the two lines (**Fig. 3D**), nearly seventy percent of genes whose promoters are bound by POU1F1 in the differentiated line are also bound by POU1F1 in the precursor line (**Fig. 3E**). We found that POU1F1 binding is associated with higher levels of gene expression in both cell lines (**Fig. 3F, 3G**).

POU1F1 binding is associated with higher ATAC-seq signal in GHF-T1 than in TαT1 cells (**Fig. 4A**). In both cell lines, POU1F1 is associated with less open chromatin than TPIT (Mayran et al. 2019). As expected sites of POU1F1 binding specific to TαT1 cells are more open in TαT1 cells (**Fig. 4B**), sites of POU1F1 binding specific to GHF-T1 cells are more open in GHF-T1 cells (**Fig. 4C**), and shared POU1F1 sites have similar signatures of open chromatin in both cell lines (**Fig. 4D**).

**Figure 4:**
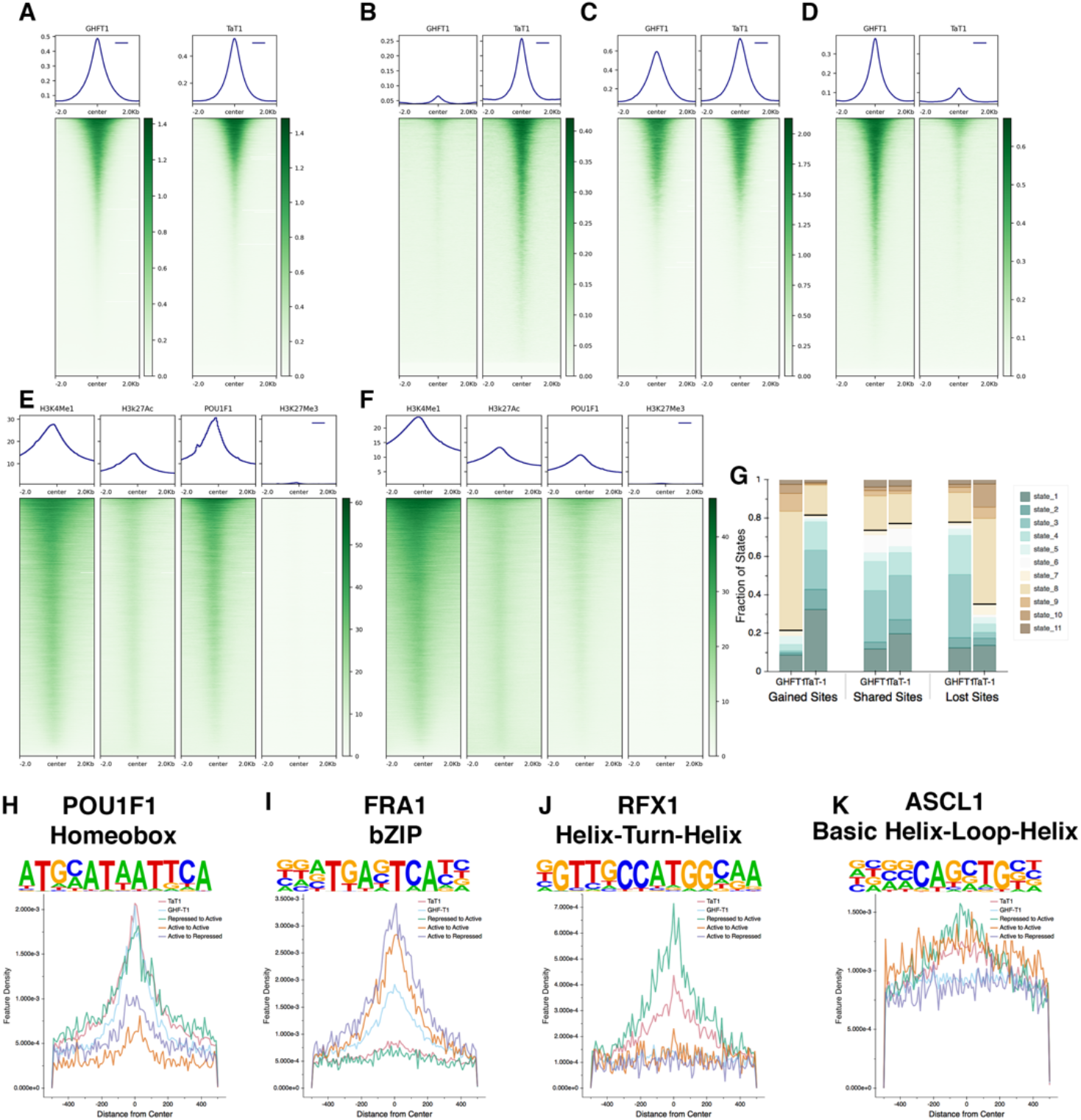
(**A**) ATAC-seq signal at POU1F1 binding sites in GHFT1 and TαT1 cells. (**B**) ATAC-seq signal at POU1F1 binding sites that are specific to TαT1 cells. (**C**) ATAC-seq signal at POU1F1 binding sites that are shared between T α T1 and GHF-T1 cells. (**D**) ATAC-seq signal at POU1F1 binding sites that are specific to GHF-T1 cells. (**E**) POU1F1 signal at enhancers in GHFT1 cells. (**F**) POU1F1 signal at enhancers in TαT1 cells. (**G**) Composition of TαT1-specific POU1F1 binding site chromatin states in GHF-T1 and TαT1 cells (gained sites, left, emissions found in **Supplemental Figure 6**), composition of shared POU1F1 binding site chromatin states in GHF-T1 and TαT1 cells (shared sites, center), composition of GHF-T1-specific POU1F1 binding site chromatin states in GHF-T1 and TαT1 cells (lost sites, right). (**H**) Density of POU1F1 motifs across POU1F1 binding sites in GHF-T1 cells (GHF-T1), TαT1 cells (TαT1), at TαT1-specific POU1F1 binding sites that are repressed in GHF-T1 cells and active in TαT1 cells (Repressed to Active), POU1F1 binding sites that are shared in GHF-T1 and TαT1 cells that are active in both (Active to Active), and POU1F1 binding sites that are specific to GHF-T1, and are in an active state in GHF-T1 cells and a repressed state in TαT1 cells. (**I**) Similar analysis as **H**, on the bZIP transcription factor, FRA1. (**J**) Similar analysis as **H**, on the HTH transcription factor, RFX1. (**K**) Similar analysis as **H**, on the bHLH transcription factor, ASCL1. **Supplemental Figure 7** shows more motifs.

Active enhancers (states containing both H3K27Ac and H3KMe1 in ChromHMM) are heavily enriched for POU1F1 binding in both cell lines (**Fig. 4E, 4F**). GHF-T1 enhancers appear to have greater POU1F1 binding than do TαT1 enhancers. POU1F1 binding that is specific to TαT1 cells is associated with active chromatin states in TαT1, but it is less so in GHF-T1 cells (**Fig. 4G**). Similarly, GHF-T1-specific POU1F1 binding is broadly associated with active chromatin states in GHF-T1 cells, but it less so in TαT1 cells. Sites bound by POU1F1 in both cell types have similarly active chromatin states, as expected.

To identify transcription factors that may be associated with differential POU1F1 binding between the cell lines, we analyzed the chromatin states associated with shared and unique POU1F1 binding sites and screened these for binding motifs. We classified genomic sites that had POU1F1 binding exclusively in TαT1 cells, and were in active states in the TαT1 cells and in repressed states in GHF-T1 cells (Repressed to Active), POU1F1 binding sites that are shared between GHF-T1 and TαT1 and are in similarly active chromatin in both (Active to Active), and GHF-T1-specific POU1F1 binding sites that are in active chromatin in GHF-T1 sites and are repressed in TαT1 cells (labeled Active to Repressed). This revealed an expected increase in POU1F1 motif density at the center of POU1F1 sites in both GHF-T1 and TαT1 cells (**Fig. 3H**). There was a striking enrichment of bZIP motifs at the center of GHF-T1-associated POU1F1 binding sites (**Fig. 3I**), suggesting that bZIP factors influence POU1F1 binding in progenitors. Interestingly, there was remarkable helix-turn-helix motif density at the center of TαT1 POU1F1 binding sites, and even more so at Repressed to Active sites (**Fig. 3J**). Similarly, there was increased bHLH motif density at Repressed to Active sites (**Fig. 3K**). These data suggest that HTH and bHLH factors mediate POU1F1 binding to novel sites in thyrotropes.

### Stretch Enhancers

Twenty-four percent of the enhancers that we identified were present in both the precursor and differentiated cell lines, as defined by at least 25% bidirectional overlap (Fig. 8A). There were 15% more enhancers in the differentiated, thyrotrope state than the precursor state. The distribution of enhancer sizes was very similar between the two cell lines (Fig. 8B). Enhancers larger than 3 kb in length, called stretch enhancers, represent 5-10% of all enhancers, are typically cell-type specific and often enriched in disease-associated areas (Parker et al. 2013; Varshney et al. 2019). Stretch enhancers represent 4.9% of the enhancer population in the precursor cell lineage and 7.1% of the enhancer population in the differentiated thyrotrope population. This is within the expected fraction, and the increased abundance in TαT1 cells is consistent with their more differentiated state. While GHF-T1 and TαT1 cells share twenty-four percent of all enhancers, only ten percent of stretch enhancers are shared between the two cell-types (**Fig. 5A**).

**Figure 5:**
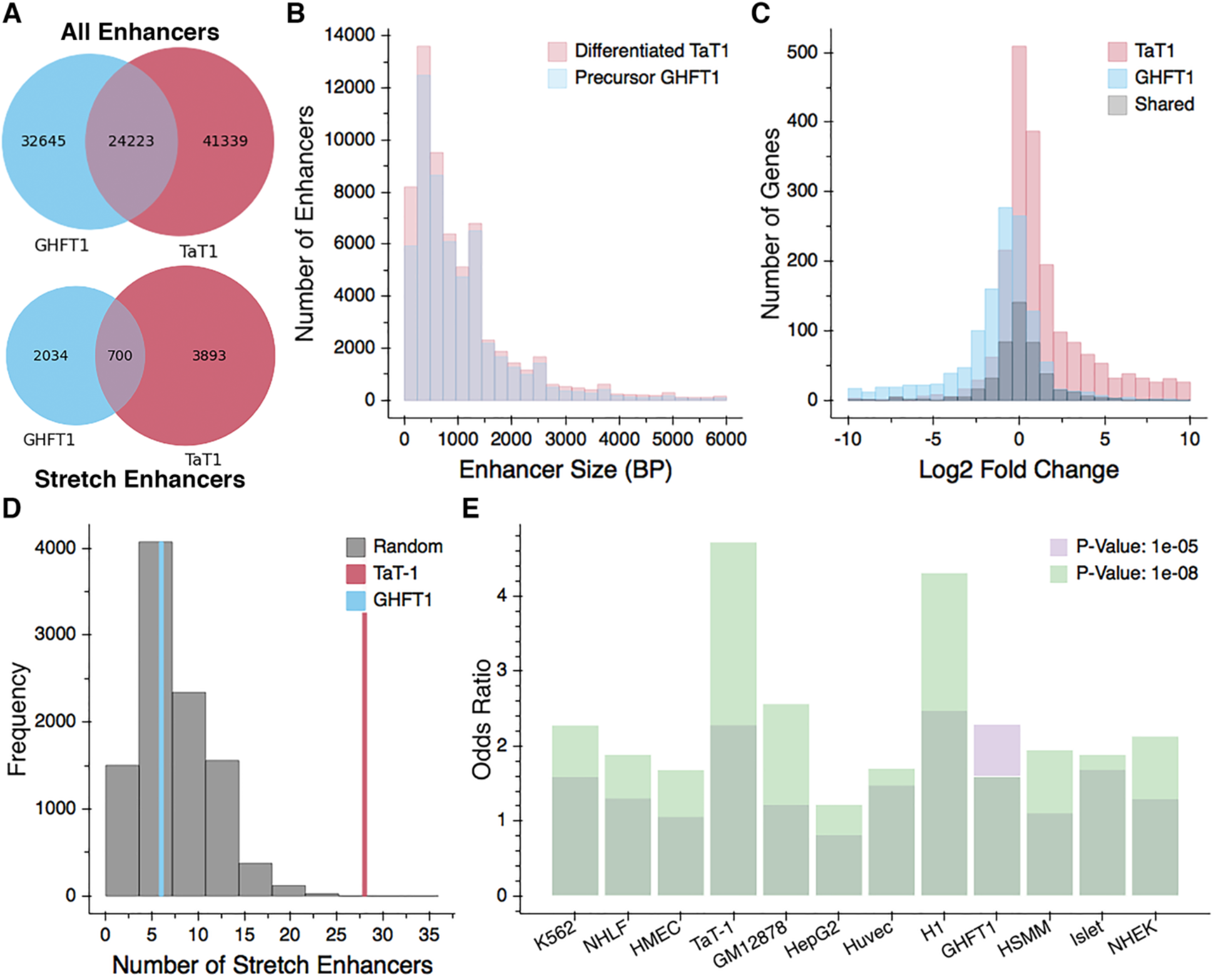
(**A**) A Venn diagram showing the number of shared and distinct enhancers in GHF-T1 and TαT1 cells (top). A Venn diagram showing the number of shared and distinct stretch enhancers in GHF-T1 and TαT1 cells (bottom). (**B**) A histogram showing the distribution of enhancer sizes in GHF-T1 (in blue) and TαT1 cells (in red). (**C**) A histogram showing the log 2-fold-change in expression of the genes that are closest to GHF-T1-specific stretch enhancers (blue), TαT1-specific stretch enhancers (red), and shared stretch enhancers (black). (**D**) Number of TαT1 (red) and GHF-T1 (blue) stretch enhancers within 100 kb (50 kb upstream or downstream) of the TSS of 25 genes important for thyrotrope function. Underneath is a histogram of the number of TαT1 stretch enhancers surrounding (within 100 kb) 10,000 iterations of 25 randomly selected genes (normalized for gene expression). (**E**) The odds ratio of observing such an enrichment of SNPs for the neuroticism subphenotype of feeling miserable within stretch enhancers of all tissues tested.

We compared the expression of the closest gene to each stretch enhancer and found that expression was highly cell-type specific. Genes closest to precursor stretch enhancers were heavily upregulated in the precursor cell line, whereas the genes closest to thyrotrope stretch enhancers were heavily upregulated in the thyrotrope cell line (**Fig. 5C**). Genes closest to shared stretch enhancers had similar gene expression in both cell lines.

We sought to determine whether genes associated with thyrotrope function in both mouse and man were closer to stretch enhancers. We generated a list of 25 candidate genes associated with thyrotrope differentiation and/or function (**Supplemental Table 3**), and we found that the TSS’s of all of these genes were within 100 kb of 28 TαT1 stretch enhancers and only 6 GHF-T1 stretch enhancers (**Fig. 5D**). To determine whether this result was significant, we randomly selected 25 genes 10,000 times, ensuring the genes had similar expression levels, and we counted the number of stretch enhancers within 100 kb of the transcription start site of those randomly selected genes. The randomly selected genes were within 100 kb of 28 TαT1 stretch enhancers only 2 times out of 10,000, yielding an empirical p-value of 0.0002, which confirms the enrichment of TαT1 stretch enhancers at thyrotrope-signature genes.

To probe the potential value of these data for application to human disease studies, we mapped the mouse enhancers in TαT1 and GHF-T1 cells from the mm9 genome onto the human genome, hg19. Because ~90% of GWAS SNPs are intronic or intergenic, and stretch enhancers are heavily enriched for disease SNPs, we expected to implicate thyrotropes in disease phenotypes by uncovering enrichment of disease SNPs in TαT1 stretch enhancers (Edwards et al. 2013; Parker et al. 2013). We used GARFIELD to measure the enrichment of these SNPs in GHF-T1 and TαT1 and stretch enhancers, while accounting for linkage disequilibrium, minor allele frequency, and distance to TSS (Iotchkova et al. 2019). We compared their enrichment to stretch enhancers found in heterologous cell lines including but not limited to Islet cells, GM12878 (human B-lymphocyte cells), and K562 (human myelogenous leukemia cells) cells (Parker et al. 2013). The study that exhibited the greatest enrichment of SNPs in TαT1 stretch enhancers (odds ratio of 4.7) was from GWAS done on the neuroticism sub-phenotype of feeling miserable (**Fig. 5E**) (Nagel et al. 2018). While the significance of this is uncertain, untreated hypothyroidism can be associated with fatigue and depression. **Supplemental Figure 8** shows the enrichment odds ratios for all GWAS studies and cell line stretch enhancers, as well as their p-values.

### *In vitro* validation of enhancers

We sought to test putative enhancers of thyrotrope-signature genes, namely *Gata2*, *Cga*, and *Pitx1*, by transient transfection of TαT1 cells. We identified putative regulatory elements as regions with significantly enriched ATAC-seq signals near these genes. The promoter proximal sequences of each gene were amplified from genomic DNA and fused to a luciferase reporter gene. Putative regulator elements were amplified from genomic DNA and cloned in both the forward and reverse orientation upstream of the promoter proximal region. The transfection efficiency of TαT1 cells is low (~20%), and the results vary, likely due to poor adherence of cells to the plate. Thus, all experiments were performed with six replicates. We detected strong enhancer elements for *Gata2* and *Pitx1* (**Fig. 6).** The position (genome coordinates) of cloned promoters and elements is presented in **Supplemental Table 4**).

**Figure 6:**
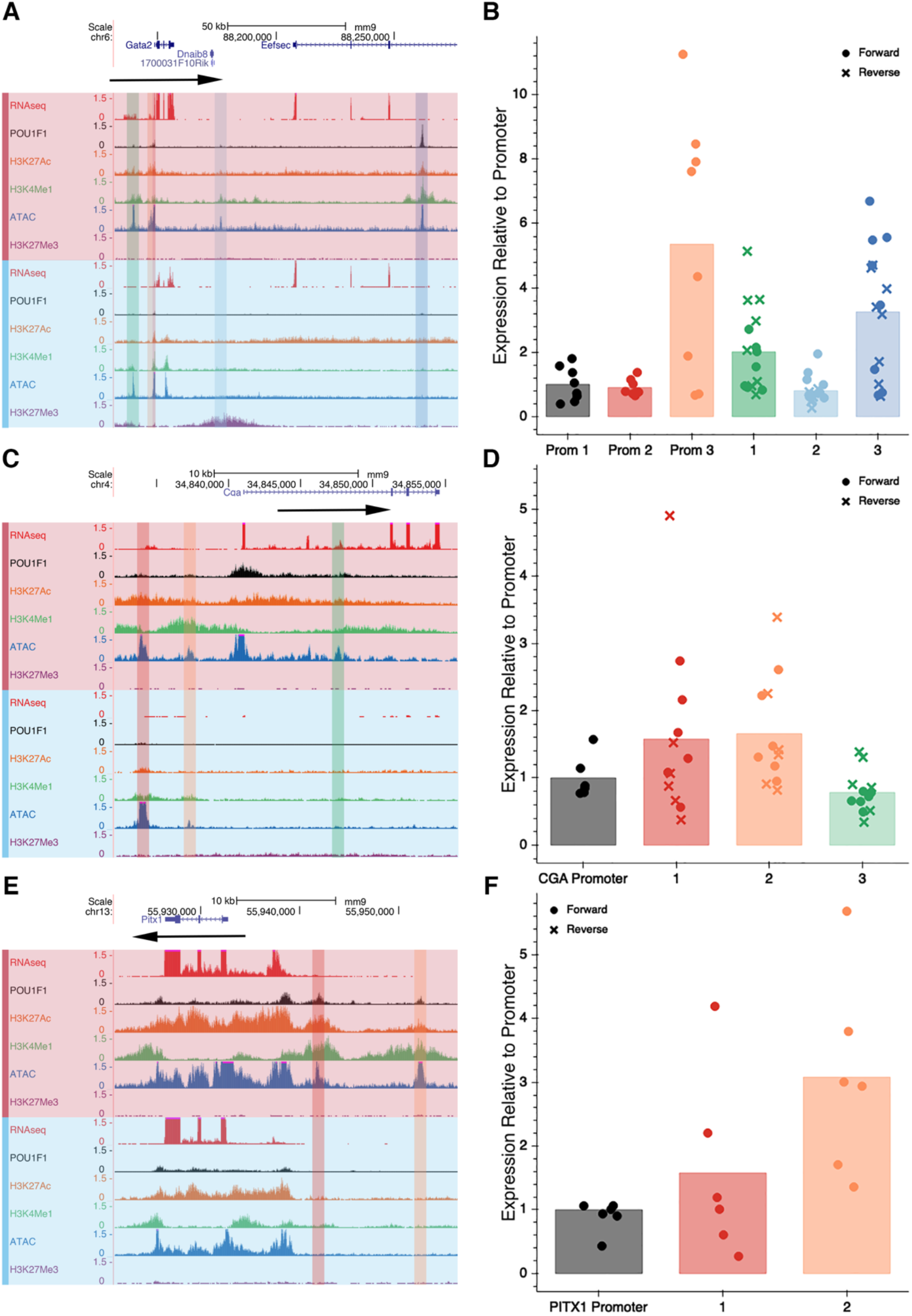
(**A**) RNA-seq, POU1F1, H3K27Ac, H3K4Me1, ATAC-seq, and H3K27Me3 tracks (TαT1 in red, GHF-T1 in blue) at *Gata2*, with the pieces of DNA cloned for the luciferase assay highlighted in red, orange, green, light blue and dark blue. (**B**) Level of luciferase activity of each element. Prom 1, 2, and 3 represent the 0.2, 0.9, and 2.8kb promoters tested, and 1, 2, and 3 represent the three similarly highlighted elements in **A** tested in both the forward (circles) and reverse (x’s) orientation upstream of the *Gata2* 0.2kb promoter. (**C**) Same tracks as in **A**, at the *Cga* locus, where elements tested are highlighted. (**D**) Level of luciferase activity of each element, color-coordinated with the highlighted elements in **C** in both the forward (circles) and reverse (x’s) orientation. (**E**) Same tracks as in **A** at the *Pitx1* locus, where elements tested are highlighted. (**F**) Level of luciferase activity of each element, color-coordinated with the highlighted elements in **F**. Elements were tested only in the forward orientation.

*Gata2* is implicated in *Tshb* transcription and proper thyrotrope function (Gordon et al. 1997; Dasen et al. 1999; Charles et al. 2006). *Gata2* has two promoters, and each are upstream of a non-coding exon (Minegishi et al. 1998). The more distal promoter is located ~5 kb upstream of the more proximal promoter, and it drives expression in Sca-1+/c-kit+ hematopoietic progenitor cells, while the downstream promoter drives *Gata2* expression in most other tissues. According to our RNA-seq data the downstream promoter is the only one utilized in TαT1 and GHF-T1 cells. *Gata2* expression is five-fold higher in TαT1 cells than in GHF-T1 cells and there is a larger area of accessible chromatin upstream of *Gata2* in the TαT1 cells (~2.8 kb vs 0.9 kb, **Fig. 6A**). There is no difference in activity between the 0.2 kb and 0.9 kb promoter proximal region in TαT1 cells, but the larger, 2.8 kb promoter-proximal region of *Gata2* stimulated luciferase activity 5-fold (p-value = 0.009). This indicates the presence of enhancer elements between 0.9 and 2.8 kb of the common TSS for *Gata2*. We tested three distal elements, fusing them with the smallest, 0.2 kb *Gata2* promoter construct. An element ~100 kb 3’ of the *Gata2* common promoter had significant POU1F1 binding and drove the highest levels of luciferase expression, (3-fold increase, p-value = 0.005). Thus, we identified two enhancer elements for *Gata2* expression in thyrotropes, one in the proximal promoter region, within 2.8 kb of the TSS, and a more distal one, approximately 110 kb downstream.

*Cga* is the alpha-subunit of TSH, and the gonadotropins, FSH and LH, and it is expressed in both thyrotropes and gonadotropes. We tested three *Cga* enhancer elements, and two appeared to have activity, although they did not reach statistical significance. The element ~7 kb upstream of *Cga* increased luciferase activity 1.5-fold, and it has not been previously described. The element located 4.6 kb upstream of the *Cga* gene increased luciferase activity 1.6-fold and approached statistical significance (p-value = 0.07). This element was previously demonstrated to be sufficient for developmental activation, cell type specific expression, and hormonal regulation in transgenic mice (**Fig. 6B**) (Brinkmeier et al. 1998; Wood et al. 1999).

*Pitx1* has a role in *Pomc* expression and hindlimb formation in mice and humans (Lamonerie et al. 1996; Daly and Camper 2020). *Pitx1* and the related *Pitx2* gene drive early pituitary development and are expressed in thyrotropes and gonadotropes (Charles et al. 2005). Mice with a pituitary-specific disruption of *Pitx2* exhibit elevated *Pitx1* expression in response to induced hypothyroidism, suggesting functional compensation (Castinetti et al. 2011). The *Pitx1* regulatory landscape extends over 400 kb and includes a pituitary enhancer 110 kb upstream (Kragesteen et al. 2018). ATAC-seq signatures revealed two previously undescribed thyrotrope-specific regions of open chromatin 9 and 19 kb upstream of the *Pitx1* TSS (**Fig. 6C**). The proximal element did not have statistically significant enhancer activity. The distal element, however, increased luciferase activity 3-fold (p-value = 0.009).

### Discovery of a novel TSH β-subunit enhancer

A bacterial artificial chromosome clone containing *Tshb* and 150 kb of surrounding DNA sequence was sufficient to drive expression in thyrotropes of transgenic mice (Castinetti et al. 2011), but there is no information about the location of key regulatory elements within this region. In fact, multiple efforts to drive expression in transgenic mice with smaller constructs were unsuccessful (Camper et al. 1995). We sought to leverage the information we have about the chromatin states in the TαT1 cells to identify elements sufficient for *Tshb* expression in mice. Knowing that the 150 kb BAC was sufficient to drive expression in thyrotropes, we limited our search to this space and found five areas with high ATAC-seq signal in TαT1 cells (**Fig. 7A**). We tested each of these elements both in the forward and reverse orientation fused to a 0.4 kb *Tshb* promoter proximal region driving luciferase expression (**Fig. 7B,** genomic coordinates of the cloned promoter and putative enhancer elements is in **Supplemental Table 4**). We discovered that an element 7 kb upstream of the *Tshb* TSS drove significant levels of luciferase (hereafter named Element 4). Element 4 had extensive ATAC-seq signal and H3K4Me1 and POU1F1 binding, consistent with the observation that POU1F1 is important for *Tshb* expression. We tested the ability of GATA2 and POU1F1 to activate *Tshb* promoter alone and in conjunction with Element 4 in heterologous CV1 cells (Gordon et al. 1997). GATA2 and POU1F1 independently cause modest increases in *Tshb*-luc reporter gene expression (2.7-fold and 1.6-fold respectively), and together they drive higher expression (5-fold, **Fig. 7C**). In the presence of Element 4, POU1F1 has very modest effects on reporter gene expression (1.3-fold increase), but GATA2 has a strong effect (4.3-fold increase). GATA2 and POU1F1 do not have an additive effect on element 4 reporter activity (4.4-fold increase) in contrast to the promoter proximal region. This suggests that GATA2 is a powerful regulator of *Tshb* expression through interaction with Element 4. To determine which other factors may be binding Element 4, we checked for the presence of over 1,000 Jaspar motifs within Element 4 at an 80% threshold. We found extensive predicted GATA2 and PITX1 binding (**Fig. 7D**). A more complete list of predicted binding factors is presented in **Supplemental Table 5**.

**Figure 7:**
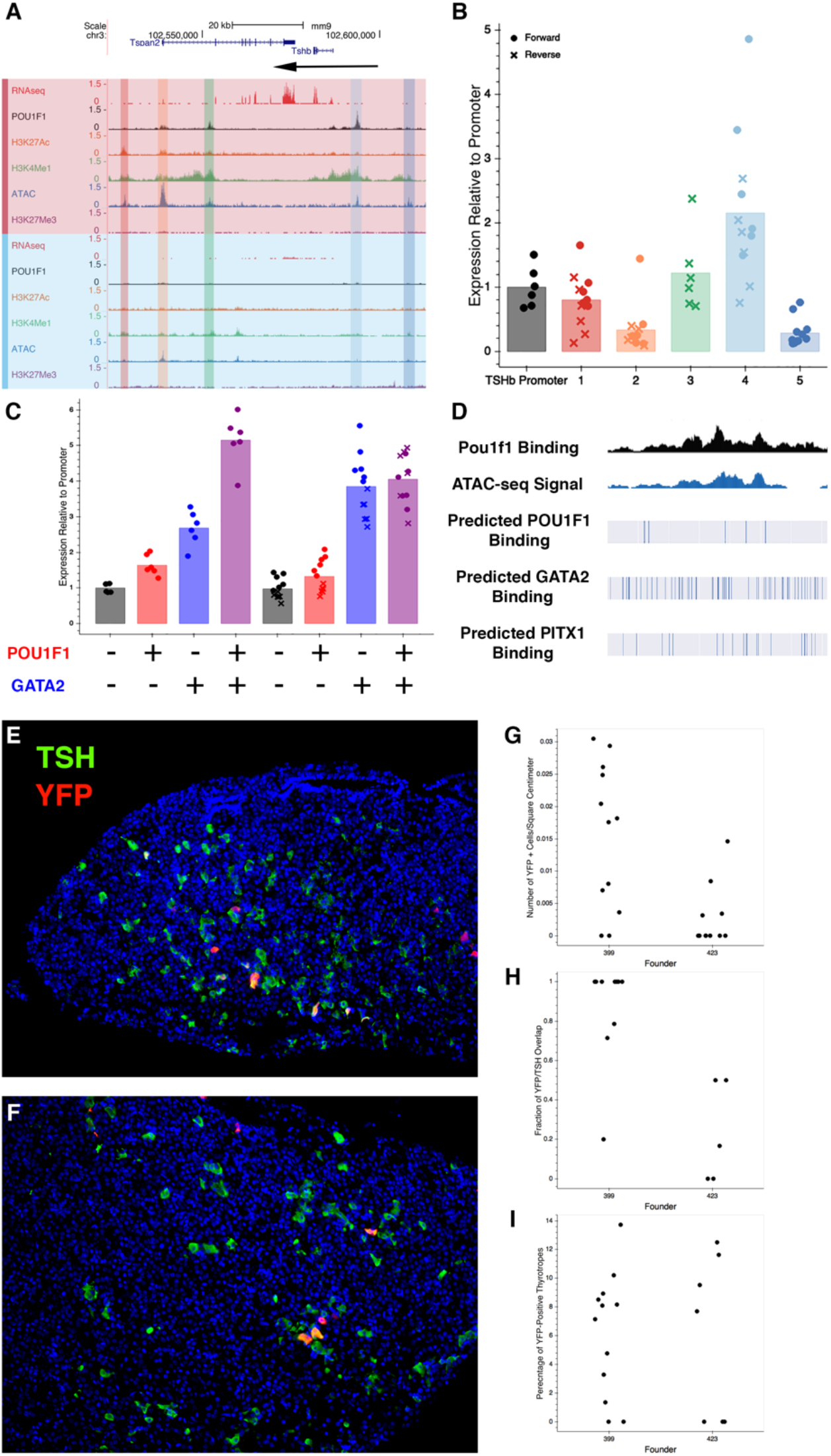
(**A**) *Tshb* locus with genome browser tracks illustrating RNA-seq, CUT&RUN for POU1F1, H3K27Ac, and H3K4Me1, ATACseq, and CUT&RUN for H3K27Me3 in TαT1 cells (red) and GHF-T1 cells (blue). Elements that were tested functionally are highlighted. (**B**) TαT1 cells were transfected with the *Tshb* promoter proximal element fused to luciferase and putative regulatory elements in the forward (filled circles) or reverse (x) orientation. Luciferase activity for each construct is normalized to the promoter-only element. (**C**) Heterologous CV1 cells were transfected with the element 4 construct and with expression vectors for POU1F1 and/or GATA2 as indicated by + or −. (**D**) Genome browser track illustrating Element 4 (1.4 kb) with experimentally determined POU1F1 binding sites and ATAC-seq sites, and predicted binding sites for POU1F1, GATA2, and PITX1 that reach a confidence level of at least 0.8 in JASPAR. An extended list of motifs is presented in Element 4 is in **Supplemental Table 5**. (**E**) Pituitary gene expression analysis in transgenic founder 399 with coimmunostaining for YFP (red) and TSHB (green), revealing overlap in expression (yellow). (**F**) Same as E, in founder mouse 423. (**G**) The number of YFP-positive cells per unit area in each founder. (**H**) The percentage of thyrotropes that express YFP in each founder. (**I**) The fraction of YFP-positive cells that are thyrotropes in each founder.

We tested whether Element 4 was sufficient to drive expression in transgenic mice by placing the Element 4 in front of the *Tshb* promoter and a YFP reporter gene. This construct was injected into fertilized eggs that were subsequently transferred to pseudopregnant foster mothers. We dissected the pituitaries of eleven founder transgenic mice at four weeks of age and examined the expression of YFP using immunohistochemistry (**Fig. 7E, 7F**). Six founders had no detectable YFP activity in the pituitary gland, three founders had low levels of YFP activity, and two founder mice had higher levels of YFP activity. We analyzed the latter two in more detail. 87% of YFP-positive cells were also positive for TSH in one founder, indicating high specificity for thyrotropes (**Fig. 7H**). 6% of the transgenic thyrotropes were also positive for YFP, suggesting low penetrance of expression (**Fig. 7I**). This represents the first regulatory element that is sufficient to drive reporter expression in thyrotropes, as several other constructs were insufficient for *in vivo* expression (Camper et al. 1995). This serves as a proof of the principle that combined transcriptome and epigenome data can be valuable for identifying enhancer elements that function in developmentally specific cell lines and intact animals.

## Discussion

Our work builds on the ENCODE effort to discover regulatory elements in diverse tissues. This represents the first systematic characterization of the epigenome and transcriptome of a thyrotrope-like cell line, providing insight into the changes that are associated with the differentiation of committed POU1F1 pituitary cells into thyrotropes. POU1F1 has a similar binding profile at promoters in the two cell lines, but there are novel binding sites at enhancers in each cell line that are associated with striking shifts in chromatin states. Global analysis of positive histone marks (H3K27Ac and H3K4Me1) revealed putative active enhancers in the two cell lines. We demonstrated that many of the enhancers surrounding thyrotrope-signature genes drive expression in a thyrotrope cell line. Furthermore, an enhancer element upstream of *Tshb* proved to be sufficient to drive expression in thyrotropes in transgenic mice. The transcriptomic, epigenomic, and POU1F1 binding data here contributes significantly to our understanding of thyrotrope cell specification, which are the key cells for regulation of thyroid gland development and growth and metabolism.

As hormone-producing cells mature, they ramp up translational machinery for robust hormone production, and CREB3L2 is a master regulator of this process in the pituitary corticotropes (Khetchoumian et al. 2019). While *Creb3l2* is not highly expressed in TαT1 cells, *Creb3l1* is expressed nearly 30-fold higher in TαT1 cells than in GHF-T1 cells (172.5 FPKM vs. 5.9 FPKM). It is possible thyrotropes use a similar mechanism of increasing translation to meet the demand for thyrotropin. Consistent with this, *Creb3l1* is upregulated in a model of thyrotrope adenoma (Gergics et al. 2016).

Pituitary endocrine cells are electrically excitable, and voltage-gated calcium influx is the major trigger for hormone secretion (Fletcher et al. 2018). G-protein coupled receptors, ion channels, and hormones all are considered components of cellular identity. For example, thyrotropes have unique electrical activity relative to other pituitary hormone-producing cell types. TαT1 cells exhibit a bursting pattern of action potentials that are affected by exposure to TRH and thyroid hormone, but the nature of the ion channels regulating TSH secretion is not understood (Mollard et al. 1990; Tomic et al. 2015). The involvement of ion channels in excitation-secretion coupling is an area of active study. The hypothalamic factors CRH, TRH, GHRH and somatostatin have an effect on electrical activity in corticotropes, lactotropes and somatotropes. Thyrotropes have not been well studied in this regard. Our study provides evidence for the acquisition of ion channel gene expression as progenitors adopt the thyrotrope fate. Voltage gated potassium channels *Kcnc3*, *Kcnq2*, *Kcnk1*, and *Kcnk2* were highly expressed in TαT1 cells. Interestingly, *KCNQ1* missense mutations cause growth hormone deficiency (Tommiska et al. 2017). We found that sodium channel genes are upregulated in two cell types that express hormones, Pit1-triple and TαT1. The calcium channel CACNA1G, a low-voltage activated, T-type channel, was highly elevated in TαT1 cells. This suggests that the previously proposed model, in which ion channel expression is pruned as differentiation proceeds, needs to be updated (Fletcher et al. 2018). Knowing which ion channels are expressed in thyrotrope cells is the first step in understanding the mechanism whereby TRH stimulates TSH release in a pulsatile manner and according to the appropriate diurnal rhythm.

Two pituitary cell types express POMC, corticotropes and melanotropes, and they process the protein differently to make adrenocorticotropin and melanocyte-stimulating hormone, respectively. Although there are very few differences in the transcriptomes of these cells, a single pioneering transcription factor, PAX7, remodels the chromatin landscape and provides access to new binding sites for TBX19 (Tpit), which drives melanotrope fate (Budry et al. 2012; Mayran et al. 2018; Mayran et al. 2019). We observed far more differences in gene expression between an undifferentiated GHF-T1 cells and differentiated TαT1 cells, however, suggesting that a single factor may not responsible for all of differences between these cell states.

Several transcription factors were strongly upregulated in TαT1 cells relative to GHFT1, including ISL1, RXRG and LHX3. Upregulation of *Isl1* and *Rxrg* was expected because pituitary-specific deletion of *Isl1* causes reduced thyrotrope differentiation (Brinkmeier et al. 2020), and several lines of evidence support a role for *Rxrg.* RXRG suppresses serum TSH levels and *Tshb* transcription, *Rxrg* deficient mice have central resistance to thyroid hormone, and loss of retinoic acid signaling suppresses thyrotrope differentiation (Brown et al. 2000; Cheung and Camper 2020). *Isl1* had extensive POU1F1 binding across the 1 MB region surrounding it. *Lhx3* expression is detectable at e9.5 in the mouse pituitary placode and expression persists though adulthood (Sheng et al. 1997). Thus, we expected to detect *Lhx3* transcripts in all pituitary cell lines. *Lhx3* transcripts were nearly undetectable in Pit1-zero cells and in the precursor GHF-T1 lineage, but transcripts were higher in Pit1-triple (1.6 FPKM) and highest in TαT1 cells (44.2 FPKM). Recently, an SV40-transformed pituitary precursor cell line was developed that expresses the stem cell marker SOX2 but not LHX3 (Daly). There may be dynamic changes in *Lhx3* expression during development that have not been documented.

SHH signaling is critical for establishing the pituitary placode and induction of *Lhx3* expression (Carreno et al. 2017). The GHFT1 precursor lineage expressed downstream targets of SHH, *Gli2* and *Gli3*, at higher levels than TαT1 cells. This is suggestive of active SHH signaling. *Gli2* and *Gli3* promoters are associated with extensive H3K4Me1 and H3K27Ac, and active enhancers can be found upstream, downstream, and within their introns. By contrast, *Gli2* and *Gli3* have broad stretches of H3K27Me3 in the TαT1 line, a mark of active repression. The active expression of these elements in GHF-T1 cells underline how well these cell types represent the early pituitary state, and suggest they could be for valuable for identifying GLI target genes that underlie pituitary developmental abnormalities (Fang et al. 2016).

POU1F1 is critical for development of thyrotropes, somatotropes and lactotropes, and it likely interacts with other factors that specify the three different cell fates. Motif analysis near unique POU1F1 sites suggests which other factors may be involved. POU1F1 binding is associated with the homeobox motif, and sites of TαT-1-specific POU1F1 binding that are repressed in GHF-T1 cells and active in TαT1 cells are heavily enriched for bHLH and HTH motifs. This raises the possibility that bHLH and HTH factors pioneer the binding of POU1F1 which then activates thyrotrope-specific expression.

Members of the RFX family of transcription factors are attractive candidates for interaction with POU1F1 to drive thyrotrope fate. These HTH factors contain DNA binding and heterodimerization domains and regulate cell fate in many organ systems, including the pancreatic islets and the sensory cells of the inner ear (Ait-Lounis et al. 2007; Elkon et al. 2015). They interact with other POU and SIX factors to direct fate. *Rfx1, Rfx2, Rfx3, Rfx5,* and *Rfx7* are expressed in both GHF-T1 and TαT1 cells. *Rfx3, Rfx4, Rfx5* and *Rfx7* are expressed in pituitary development between e12.5 and e14.5, a time when progenitors leave the cell cycle and initiate differentiation (Brinkmeier et al. 2009). Future studies will be necessary to define the role of these genes in pituitary development.

The expression of bHLH factor genes *Ascl1* (*Mash1*), and *NeuroD4* (*Math3*), are elevated in TαT1 cells. However, thyrotrope commitment is normal in *Ascl1* knockout and in triple knockout mice deficient in *Ascl1, Neurod4,* and *Neurod1.* These bHLH activating factors have overlapping functions in promoting somatotrope, gonadotrope, and corticotrope development (Herzog et al. 2004; Pogoda et al. 2006; Zhang et al. 2015; Ando et al. 2018). They promote pituitary stem cell exit from the cell cycle and act as selectors of cell fate, as triple knockout mice have more SOX2-positive cells and more lactotropes. bHLH factors play important roles in neuronal differentiation. For example, the BAM factors ASCL1, BRN2 and MYT1L are sufficient to transdifferentiate mouse embryonic fibroblasts (MEFs) into neurons. The Zn finger transcription factor MYT1L is highly upregulated in TαT1 cells, and although the POU factor *Brn2* is barely expressed in TαT1 cells, POU1F1 is highly expressed. Thus, it is possible that ASCL1 acts in concert with MYT1L and POU1F1 to drive thyrotrope fate.

The top three most highly expressed bHLH factors in both of pituitary cell lines are the repressive bHLH factors in the ID family. The role of these genes in pituitary development has not been studied, but they may be important in regulating progenitor differentiation and cell fate selection. For example, a proper balance of activating and repressive bHLH factors is critical for cortical development (Ross et al. 2003). Repressive ID and Hes factors are expressed in cortical progenitors, and induction of key activating bHLH factors drive these progenitors to differentiate. Astrocytes, however, require continued repressive bHLH factor expression. The interplay of active and repressive bHLH factors in pituitary development is likely complex.

This work represents the first thorough characterization of the epigenome and transcriptome in *Pou1f1* lineage progenitors and thyrotropes. We used this genome-wide catalog and tested enhancer function of elements in genes encoding thyrotropin, *Cga* and *Tshb*, the receptor for the hypothalamic releasing hormone that regulates thyrotropin, *Trhr*, and two crucial transcription factors, *Gata2* and *Pitx1*. This provides proof of the principle that the catalog is valuable for dissecting gene regulation. In addition, we demonstrate that the *Tshb* enhancer element is sufficient for expression in thyrotropes in transgenic mice and is directly regulated by GATA2. We discovered that unique POU1F1 binding sites are associated with bZIP factor binding motifs in *Pou1f1* lineage progenitors and bHLH or bHTH binding motifs in thyrotropes. This suggests candidate gene families for regulating thyrotrope differentiation, such as the RFX family. Of the more than 30 known genes that are mutated in patients with hypopituitarism, 18 are transcription factors that regulate pituitary development and cell specification, indicating the clinical importance of this field of study. The overwhelming majority of patients with hypopituitarism have no molecular diagnosis, suggesting additional genes remain to be discovered. We provide a rich selection of candidate transcription factors that are differentially expressed in progenitors and thyrotropes. Amongst the top 40 of these, 9 are already implicated in pituitary development and disease. Future analysis of the remaining 31 candidates may uncover additional disease genes.

## Methods

### Cell culture and transfection

GHF-T1 and TαT1 cells were provided by Dr. Pamela Mellon at University of California San Diego and grown on uncoated 100mm dishes and Matrigel-coated 60mm dishes, respectively. They were grown in DMEM (Gibco, 11995-065) with 10% Fetal Bovine Serum (Corning, 35016CV) and 1% Penicillin Streptomycin (Sigma-Aldrich P4333). Cells were split 1:10 once they achieved 80% confluence. Six replicates of TαT1 cells were transfected using FuGENE 6 with a 3:1 transfection reagent/DNA ratio. Cells were collected 48 hours post-transfection for collection and luciferase measurement was performed using Promega Dual-Glo (Promega #E2920), and a GloMax 96 microplate luminometer.

### Cloning

The DNA prepared for the plasmids used in the transfection experiments and transgenic mice was amplified from TαT1 DNA. Primers were designed to work with Phusion Green Hot Start II High-Fidelity PCR Master Mix (catalog # F566S). Two methods were employed to clone plasmids containing the regulatory element, the respective promoter, and the YFP reporter. The first method involved adding 10-15 nucleotides onto the insert that overlapped with the pCDNA3-YFP Basic plasmid that had been cut with Kpn1 and Xho1. The amplicon and linearized plasmid backbone were combined using the NEB HiFi DNA Assembly Master Mix (catalog # E2621L) with 1:2 vector:insert ratios, and 60 min incubation time at 50°C. The subsequent plasmid was transformed into DH5αcells. The second method was to insert the amplified construct into a Zero Blunt TOPO vector (ThermoFisher #450245), select clones that are in the forward and reverse orientation, then cut the TOPO vector containing the insert with Kpn1 and Xho1. The resulting fragment is then ligated into a digested pCDNA3-YFP Basic plasmid that had been cut with Kpn1 and Xho1 using a standard T4 DNA Ligase protocol (NEB #M0202). Once the insert, plasmid, and breakpoint sequences were confirmed by Sanger sequencing, the plasmids were extracted from overnight 1 L cultures of DH5αcells using Qiagen Plasmid Maxi Kits (catalog #12163). Endotoxins were removed from the plasmids using the Endotoxin removal Solution (Sigma, E4274-25ML).

### RNA Seq

One million GHF-T1 and TαT1 cells were collected for each of the three replicates for each cell line. Once collected, the RNA was extracted using the RNAqueous™ Total RNA Isolation Kit (catalog #AM1912). The RNA was prepared by the University of Michigan Advanced Genomic Core for mRNA enrichment followed by 50-cycle, paired-end sequencing on the Illumina HiSeq-4000. The RNA was checked for quality using FastQC and mapped and analyzed using the VIPER Snakemake pipeline (Cornwell et al. 2018). Briefly, VIPER aligns the files to the mm9 transcriptome using STAR, followed by differential expression analysis using DESeq2 and celltype clustering and expression quantification using QoRTs (Hartley and Mullikin 2015). The quality of the alignment was also analyzed using QoRTs.

We measured the significance of the increase in expression of sodium, potassium, and calcium using a one-way ANOVA. This demonstrated the significance of the increase in expression of the sodium and potassium, but not the calcium channel genes.

### GO Term and KEGG Pathway Enrichment

We performed GO-term enrichment on the top 5% of most differentially expressed genes (by log-2 fold-change) in both lines (Ashburner et al. 2000; The Gene Ontology 2019). This represented 453 genes in GHFT1 cells, and 490 genes in TαT1 cells. We used the default settings on the web-based Gene Ontology Resource (geneontology.org), using the biological process and Mus Musculus options. The resulting GO terms were plotted by their log-2 fold enrichment, and their p-values.

We also performed a directional Kyoto Encyclopedia of Genes and Genomes (KEGG) pathway enrichment analysis on all of the genes using RNA-enrich (Lee et al. 2016). We set the maximum number of genes per concept to 500 and the minimum number of genes per concept to 5, and otherwise used the default settings. The resulting KEGG pathways were plotted by their coefficients and p-values.

### ATAC-Seq

The Assay for Transposase Accessible Chromatin with high-throughput sequencing (ATAC-seq) was performed as previously described (Buenrostro et al. 2013; Buenrostro et al. 2015). Briefly, 50,000 nuclei were extracted from collected GHF-T1 and TαT1 cells. The cells were transposed with Illumina transposase (Illumina #FC-121-1030) for 30 minutes at 37°C while shaking at 250 RPM. The resulting fragmented DNA was amplified using ¼ of the cycles required to reach saturation in the described qPCR QC. The final amplified DNA library was purified using the Qiagen PCR purification kit (catalog #28104) and sequenced on the Illumina HiSeq platform. The quality of the reads was checked using FastQC, aligned to the mm9 genome, and had its peaks called using the Parker lab’s Snakemake pipeline (Rai et al. 2020).

### CUT&RUN

CUT&RUN was performed under high-digitonin conditions as described with few exceptions, namely all steps with < 1 ml of liquid requiring mixing were done by 500 RPM shaking instead of inversion (Skene and Henikoff 2017). Briefly, 250,000 cells per sample, and one sample per cell line-antibody pair were collected, washed, and bound to Concavalin A beads (Bangs Laboratories, BP531). The cells attached to beads were incubated at 4°C overnight with the respective antibodies (POU1F1 – this antibody was a kind gift from Dr. Simon Rhodes, University of North Florida, Jacksonville, FL (Prince et al. 2013), H3K27Me3 – Cell Signal #C36B11, H3K27Ac – Abcam ab4729, H3K4Me1 – Abcam ab8895, Rabbit IGG – R&D AB-105-C). The antibodies were washed, and no secondary antibody was used. The protein A/MNAse fusion protein was added, followed by Ca^2+^-induced digestion at 0°C for 30 minutes. The fragmented chromatin was then collected and purified using Macherey-Nagel NucleoSpin Gel and cleanup columns (catalog #740609). Libraries of this DNA was prepared using the Kapa Biosystems library prep kit (catalog #KK8702) at a 100:1 adapter:sample ratio. The libraries were paired-end sequenced on a single lane of the Illumina HiSeq-4000 for 50 cycles. The resulting data was then checked for quality using FastQC, aligned to the mm9 genome using Bowtie2 using the flags recommended for CUT&RUN (--local --very-sensitive-local --no-unal --no-mixed --no-discordant --phred33 -I 10 -X 700), and peaks were called using MACS2.

### ChromHMM

ChromHMM was performed on both cell lines using H3K4Me1, H3K27Ac, ATAC-seq and H3K27Me3 as input. The number of states was iteratively increased to find the number of states that resulted in the fewest number of states with the best log-likelihood. An eleven-state model was selected as a result. Association of each state with the various marks and genomic features can be seen in **Supplementary Figure 6**. Contiguous states of value two and three were stitched together as enhancers using bedtool’s mergeBed function with a -d 1 flag (Quinlan and Hall 2010).

### Association Enrichment Test

Enrichment analysis of disease SNPs at stretch enhancers in GHFT1, TαT1 and heterologous cell lines was performed using GARFIELD and stretch enhancers published previously (Parker et al. 2013; Iotchkova et al. 2019). To find the human sequences orthologous to GHFT1 and TαT1 stretch enhancers we used a conversion file generated using bnMapper and an mm9 to hg19 chain file (Orchard et al. 2019). We selected associations that had association counts of at least 50, that had a full complement of summary statistics, had more than three million tested SNPs, and were in the harmonized data format were chosen from the GWAS catalog (Buniello et al. 2019). The resulting heatmap of odds ratios and p-values for each association, tissue-type pair is shown in **Supplementary Figure 8**.

### Transgenic mice

All mice were housed in a 12-h light, −12 h dark cycle in ventilated cates with unlimited access to tap water and Purina 5020 chow. All procedures were conducted in accordance with the principles and procedures outlined in the National Institutes of Health Guidelines on the Care and Use of Experimental Animals and approved by our Institutional Animal Care and Use Committee.

Recombinant DNA was generated by amplifying genomic mouse DNA regions in **Supplemental Table 4**, and the previously described Phusion polymerase. The putative regulatory element was then combined with 438 base pairs of the TSHβpromoter (chr3:102,586,594-102,587,032), and a YFP reporter. These elements were combined using the DNA Hifi reaction into a pGEM-T Easy plasmid. The putative regulatory element, the promoter, the YFP, and the breakpoints were checked for accuracy with Sanger sequencing. Once the plasmid was confirmed, larger quantities of the plasmid were generated from overnight 1L cultures of DH5αcells using Qiagen Plasmid Maxi Kits (catalog #12163). To reduce the effect of the plasmid backbone on the viability of injected eggs, the enhancer, promoter, and YFP were amplified from the plasmid. The resulting amplicon was gel purified and injected into fertilized eggs of mice on a C57BL/6 and SJL mixed background. The resulting mice were genotyped for the YFP allele according to the Jackson Laboratory recommended primers and conditions (Srinivas et al. 2001). We dissected the pituitaries from mice that were positive for the YFP transgene (and four age-matched, negative, control littermates) at three weeks of age.

### Tissue Preparation and Immunohistochemistry

Mouse pituitaries were fixed in 4% formaldehyde in PBS overnight at 4°C. The tissue was washed three times in PBS and put in 10% EDTA for 3 hours. They were then dehydrated by putting them in 25%, then 50%, then 70% ethanol for one hour each. The pituitaries were embedded in paraffin with four-hour cycles using a Tissue Tek VIP Paraffin tissue processing machine (Miles Scientific). The embedded pituitary was cut into coronal, six-micron sections, and was analyzed by immunohistochemical markers as previously described (Castinetti et al. 2011; Mortensen et al. 2015). Anti-YFP and anti-*Tshb* were used (from Abcam ab6556 and National Hormone and Peptide Program, respectively).

Antibodies were detected using either the tyramide signal amplification (TSA) (33002 CFF488A Streptavidin HRP, Biotium, Fremont, CA) and streptavidin-conjugated Alexa-fluor 488 (1 : 200, S11223, Invitrogen). DAPI (1:200) was incubated on the slides for five minutes to stain nuclei. DABCO-containing permount was used to mount the slides, which were then imaged using a Leica DMRB fluorescent microscope.

## Data Access

All raw sequencing data generated in this study have been submitted to the NCBI Sequence Read Archive (SRA, https://www.ncbi.nlm.nih.gov/sra) under accession number PRJNA643917, and will be released once published. The reviewer link can be found here.

## Funding

NIH R01HD034283 (SAC), NIH T32GM007544 and T32HG000040 (AZD). We thank Wanda Filipak and Galina Gavrilina of the transgenic animal model core for assistance with the transgenic reporter mice (NIH P30CA046592). We thank Alex Mayran and Jacques Drouin for the advice and pituitaries of *Ascl1*-null mice. We thank Amanda Mortensen for the genotyping of the founder transgenic reporter mice. We thank Peter Orchard, Arushi Varshney, and Ricardo Albanus for their contributions to the work.

## Author contributions

Alexandre Z. Daly performed the RNA-seq, ATAC-seq, and CUT&RUN (for POU1F1, H3K27Ac, H3K4Me1, and H3K27Me3) library preparation and data analysis. He generated the plasmids and performed the transfections for the luciferase assays. He dissected the founder reporter mice, processed their pituitaries, and performed immunohistochemistry on them and processed the resulting data. He also wrote the manuscript. Lindsey A. Dudley performed immunohistochemistry on the *Ascl1*-null mouse pituitaries. Michael T. Peel and Stephen A. Liebhaber contributed RNA-seq data from Pit1-Zero and Pit1-Triple cell lines. Stephen C. J. Parker contributed to the analysis of the high-throughput sequencing data. Sally A. Camper led the experimental design, and wrote the manuscript.

## Disclosure Declaration

The authors have nothing to disclose.

